# Comparative Analysis of Tickling and Conspecific Play in Tame Mice and Golden Hamsters

**DOI:** 10.1101/2024.03.19.585680

**Authors:** Sarah Dagher, Darcie DeAngelo, Ren Y. Sato, Hiroaki Norimoto, Tsuyoshi Koide, Shimpei Ishiyama

## Abstract

Social play behavior is a fundamental aspect of animal interaction, shaping social bonds and enhancing cognitive capacity. While studies on human-animal play interactions have primarily focused on a few selected species, research on rodents beyond rats remains scarce. We, therefore, addressed the dynamics of social play in tame mice, selectively bred to approach human hands, and golden hamsters, comparing their responses during interactions with humans and conspecifics. Tame mice exhibited heightened playfulness with humans, marked by increased vocalizations and chasing behavior, in addition to increased interactions with tame conspecifics compared to unselected control mice. Hamsters demonstrated a stronger inclination towards conspecific interactions. Notably, vocalization patterns varied between heterospecific and conspecific engagement in both species, suggesting context-dependent communication. These findings offer insights into the evolutionary basis influencing social play across species with differing social structures. Understanding these mechanisms enriches our comprehension of the diverse pathways through which animals form social bonds.

## Introduction

Social play behavior is widely observed across numerous social species including dogs [1], cats [2, 3], birds [4] and rats [5, 6], serving as a vital mechanism for refining physical, cognitive, and social skills essential for navigating complex ecological landscapes [7]. In social mammals, such behavior is crucial for fostering the development of social bonds, communication proficiency, and hierarchical structures, thereby facilitating cooperative interactions within groups [8-10]. In rodents, particularly rats and mice, the manifestation of social play introduces a unique dimension to our understanding of this behavior. Rodents exhibit complex social structures and behavioral repertoires. For example, rats engage in intricate motor movements during rough and tumble play [5].

In social play, reward and positive reinforcement mechanisms play a crucial role, involving the mesolimbic dopaminergic system [8, 11-13]. Furthermore, pleasure and fostering social bonds through positive emotional associations are mediated by endorphins [14]. Brain structures, such as the prefrontal cortex, amygdala, and nucleus accumbens, orchestrate these neurobiological processes, shaping the reinforcing and bonding aspects of social play [6, 15-17].

Beyond conspecific interactions, playful behaviors extend to inter-species engagements. Human-animal social play is well-documented in domesticated animals [18-21]. Tickling is a unique form of social touch inducing laughter. Tickling animals by humans i.e. heterospecific play has been reported in non-human primates [22] and rats [23] so far, despite anecdotal observations of seemingly ticklish behaviors in other companion animals. Tickling induces appetitive ultrasonic vocalizations and playful chasing in rats [24-28] and is rewarding through the dopaminergic system [29]. Among researchers studying rodent social play, there is consensus that standard laboratory mice generally do not exhibit such responses to tickling, unlike rats. However, this observation lacks substantial documentation even as negative data in published literature. Previous research described selective breeding of wild mouse species for tameness, characterized by their approach towards and reduced avoidance of human hands, termed active and passive tameness, respectively [30-32]. Within 12 generations, selectively bred groups demonstrated significantly higher active tameness compared to unselected control groups [30]. These tame mice share upregulated genes in 2 genomic sequences of chromosome 11 in common with domesticated dogs. Despite these advances, whether tame mice engage in playful interactions with humans remains unexplored.

We wondered whether tame mice, similar to rats, would exhibit playful responses to tickling through appetitive vocalizations and hand chasing. Investigation of tickling response in tame mice might contribute to understanding the genetic underpinnings of playful traits and their role in shaping human-animal relationships. Furthermore, we examined how golden hamsters vocalize in response to tickling compared to their interactions with conspecifics. While conspecific play behavior in hamsters has been extensively documented [33], their interactions with humans, despite being common pets, remain largely unexplored in the literature.

We show that selective breeding for tameness in mice enhanced human-directed play, assessed by increased vocalizations and hand chasing, while hamsters exhibited a stronger preference for conspecific interactions over human interactions. Notably, vocalizations in both species were context-dependent, with distinct features observed between human-directed and conspecific play. These findings offer a unique opportunity to explore the evolutionary origins of playful behaviors in species with varying social structures and niches, enriching our understanding of the evolutionary trajectories of sociality and playfulness.

## Materials and Methods

### Animals

Nine 3-week-old male *Mus musculus* mice (comprising four tame mice and five unselected controls originating from the same stock without selection for tameness, with one control recruited on experiment day 2 following the death of another) and eight 3-week-old male *Mesocricetus auratus* golden hamsters were sourced from the National Institute of Genetics, Mishima [31], and Hokkaido University Medical Center, respectively. The animals were housed individually, with hamsters kept at Hokkaido University Medical Center and mice at the National Institute of Genetics, Mishima. They were maintained under a 12:12 h inverted light/dark cycle and had unrestricted access to food and water. Experimental procedures adhered to Japanese guidelines on animal welfare, overseen by local ethics committees, and were conducted in accordance with the animal experimentation permits, namely Permit No. R4-14 for mice and Permit No. 21-0092 for hamsters.

### Experimental setup

For the experiments involving mice, the setup comprised a gray polyvinylchloride box measuring 40 × 40 × 40 cm, utilized for both tickling and conspecific interaction tests. Throughout these tests, the environment was maintained at a subdued illumination of 20 lx, measured at the center of the floor. The open field test took place in a white polyvinylchloride box measuring 60 × 60 × 40 cm. For the experiments involving hamsters, the setup comprised a glass box measuring 30 × 30 × 30 cm with a subdued illumination of 8 lux, measured at the center of the floor, utilized for both tickling and conspecific interaction tests. The open field test for hamsters took place in a glass box measuring 45 × 45 × 30 cm with an illumination of 8 lux (center). Video recordings were captured using a video camera HDR-CX680 (Sony corporation, Tokyo, Japan) for mice experiments and a webcam (C922n, Logicool, Lausanne) for hamster experiments, both set at 30 frames per second (fps). Cameras were positioned overhead the arenas. Vocalizations were recorded using condenser ultrasound microphones (CM16/CMPA, Avisoft Bioacoustics, Berlin, Germany) positioned on top of the behavioral box at a sampling rate of 256 kHz and a 16-bit resolution using Avisoft-RECORDER USGH software (Avisoft Bioacoustics, Berlin, Germany). Synchronization of audio and video recordings was achieved using TTL pulses, as previously described [34]. The setup for mice is illustrated in Figure 1a.

**Figure 1:**
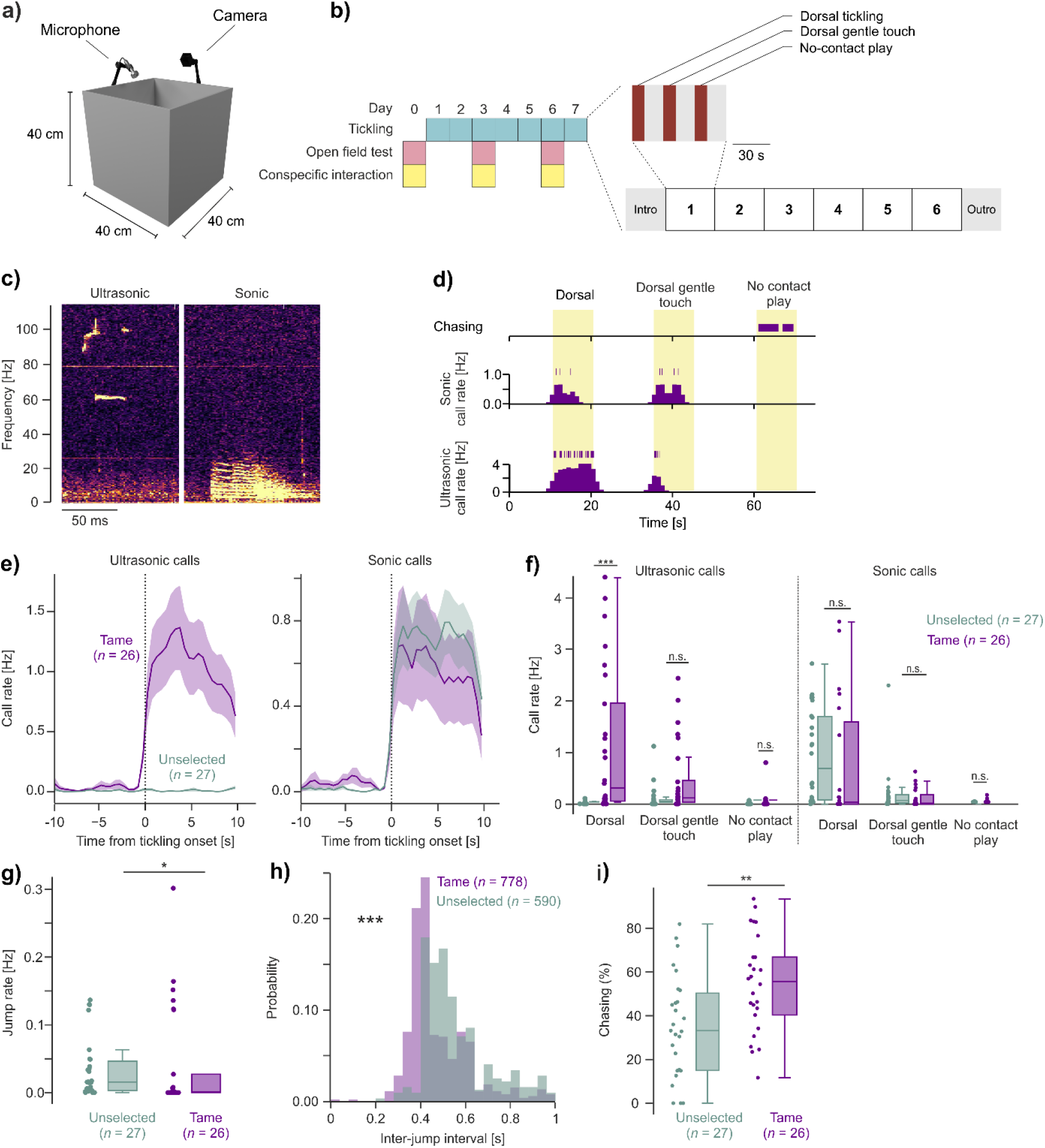
Behavioral characteristics during human-mice play interaction. **a)** Tickle arena with simultaneous audio and video recording. **b)** Left, experimental schedule consisting of tickling, open field test, and conspecific interaction test. Right, behavioral paradigm of tickling: One tickling session is a combination of 6 repetitions, each consisting of heterospecific interaction events separated by break periods. **c)** Representative spectrograms of ultrasonic (left) and sonic (right) call during tickling. **d)** Raster plot and histogram of chasing, sonic and ultrasonic call rate in a representative tickling session. **e)** Peristimulus time histogram (PSTH) of ultrasonic and sonic vocalization rate aligned to the onset of tickling. *n*: number of sessions; mean ± SEM. Data were obtained from 4 tame (purple) and 4 unselected (green) animals and binned to 500 ms. **f)** Ultrasonic and sonic vocalization rate during different heterospecific interaction events. *n*: number of sessions; rank-sum test. **g)** Jump rate during tickling sessions. *n*: number of sessions; rank-sum test. **h)** Inter-jump interval histogram. *n*: number of jumps; rank-sum test. **i)** Chasing time percentage during no-contact play phases. *n*: number of sessions; rank-sum test. All *p*-values are detailed in Supplementary Table 2.

### Experimental paradigm

#### Tickling

Tickling experiments were conducted daily from Day 1 to Day 7 for mice and from Day 1 to Day 10 for hamsters. Prior to each tickling session, animals underwent a 10 min handling period. The experimental paradigm consisted of a 2 min introductory period, followed by six repetitions of heterospecific interaction phases, namely dorsal tickling, dorsal gentle touch, and a no-contact play phase where the hand was introduced into the arena without making physical contact, allowing the animal to engage in chasing behavior. Each phase lasted for 15 s and was followed by a 10 s break. After the final repetition, animals were kept in the box for a 2 min concluding period (outro) (Figure 1b). The experimental paradigm was programmed using [Seconds Interval Timer mobile app](https://play.google.com/store/apps/details?id=com.runloop.seconds.free&pcampaignid=web_share) and remained visible to the experimenter throughout the sessions. Video recording and calls were consistently recorded during all sessions.

#### Conspecific interaction test

The conspecific interaction test was carried out on Day 0, Day 3, and Day 6 for mice, with an additional Day 9 for hamsters. To prevent any familiarity effects during the experiment, a new conspecific was introduced to each animal for every session. In the case of mice, pairs were selected from the same group, either unselected control or tame. The pairs were placed in a box for a 10 min period, during which they were allowed to interact freely, with video and calls recorded throughout the session.

#### Open field test

The open field test was conducted on Day 0, Day 3, and Day 6 for mice, with an additional Day 9 for hamsters. During each session, animals were placed in a box and given the freedom to move across the arena for a duration of 10 min.

### Analysis

#### Video analysis

Video footage was analysed using BORIS software [35] to label timings of the experimental paradigm phases as well as animal behaviors. Ethograms used for tickling and conspecific interaction video analysis are shown in Supplementary Table 1.

#### Call analysis

Call analysis flow is depicted in Supplementary Figure 1. Audio files were analysed using DeepSqueak [36] for detection of calls in both mice and hamsters. The detection results were subsequently visually cross-verified and corrected by experimenters. For mice calls (Supplementary Figure 1a), sonic and ultrasonic calls were manually assigned on DeepSqueak, and sonic calls were further confirmed in Audacity (https://www.audacityteam.org/download/) to avoid any false positives. For conspecific interaction test analysis, emitters were not identified. Features of spectrogram contour for each call including principal frequency, minimum frequency, maximum frequency, frequency standard deviation, sinuosity, mean power, tonality, peak frequency, and call duration were exported from DeepSqueak. The features were then projected to UMAP space. Difference between tickling calls and conspecific calls for both ultrasonic and sonic calls was assessed by calculating the Mahalanobis distance, incorporating bootstrapping with 1000 iterations to obtain the mean distance and its 95% confidence interval. Furthermore, each feature was compared between tickling calls and conspecific calls for both ultrasonic and sonic calls. Since manual assignment of ultrasonic and sonic calls in hamsters was not as clear as that in mice, spectrogram images of all calls were embedded into UMAP space using algorithms reported by Jourjine et al. [37] and clustered using HDBSCAN (Supplementary Figure 1b). Features of spectrogram contour extracted from DeepSqueak were compared between clusters.

#### Open field test analysis

The analysis of open field test results was performed using tracking systems, TimeOFCR1 (O’Hara & Co., Ltd., Tokyo, Japan) and Toxtrac [38] for mice and hamster files respectively.

The arena was segmented into nine equal squares by dividing it both horizontally and vertically into thirds. The central square, constituting 1/9^th^ of the total area, was designated as the center region. Total distance travelled and time spent in the center region of the arena during each session was analysed for each animal.

#### Statistical analysis

Intergroup comparisons were performed with rank-sum test for unpaired data, and Wilcoxon signed-rank test for paired data. One-sample t-test was used across various behavioral metrics to determine whether their mean values significantly differed from zero. Call cluster fraction in different events was assessed using Kruskal-Wallis test. For open field test statistical assessment and hamster call rate comparisons across events, linear mixed effect model was used with subjects as a random effect. *n* refers to sample size. Data were analysed using Python 3.8. All *p*-values for each figure are presented in Supplementary Table 2.

## Results

### Tame mice respond to tickling with ultrasonic vocalizations and played with humans

During tickle sessions involving mice, we observed both ultrasonic and sonic calls (Figure 1c). The call rates, however, varied depending on the type of interaction with humans (Figure 1d). Dorsal tickling induced ultrasonic call emissions in tame mice (Supplementary Video 1), but not in unselected control mice (Figure 1e, left; Supplementary Video 2). The increase in sonic call rate, on the other hand, was noticeable right at the onset of tickling for both tame and unselected mice, with no significant difference between the two groups (Figure 1e, right). The discrepancy in ultrasonic vocalization rate between tame and unselected mice was specifically prominent during dorsal tickling (0.275 [0.020, 2.182] vs. 0.0 [0.000, 0.009]; median & IQR; *p* < 0.001), with no significant differences observed during dorsal gentle touch and no-contact play (Figure 1f). Sonic call rates, in contrast, showed no significant differences between the groups in all interactions.

Another distinctive behavioral trait of tame mice was a significantly lower jump rate compared to unselected mice (Figure 1g; Supplementary Video 3). Tame mice exhibited jumps at shorter intervals in comparison to unselected control mice (Figure 1h). During the no-contact play phase, where mice were allowed to chase after the experimenter’s hand, tame mice displayed pronounced chasing behavior compared to unselected mice (55.7 [40.3, 66.8] vs. 33.2 [15.0, 50.4] Hz, median & IQR, *p*-value = 0.005; Figure 1i; Supplementary Video 4&5). These results demonstrate the impact of selective breeding for tameness on specific behavioral traits, particularly highlighting increased playfulness with humans.

### Tame mice engage in pronounced conspecific play with context dependent vocalizations

We previously showed that tame mice exhibit higher social preference to caged conspecifics [32]. However, their interaction with conspecifics in a freely-moving setting and its comparison to human interaction remains unknown. Therefore, we performed the conspecific test, where two mice from each group were paired together on each occasion. Tame mice pairs notably exhibited a higher frequency of interactions associated with ultrasonic vocalizations compared to unselected pairs (Figure 2a, b & Supplementary Table 1).

**Figure 2:**
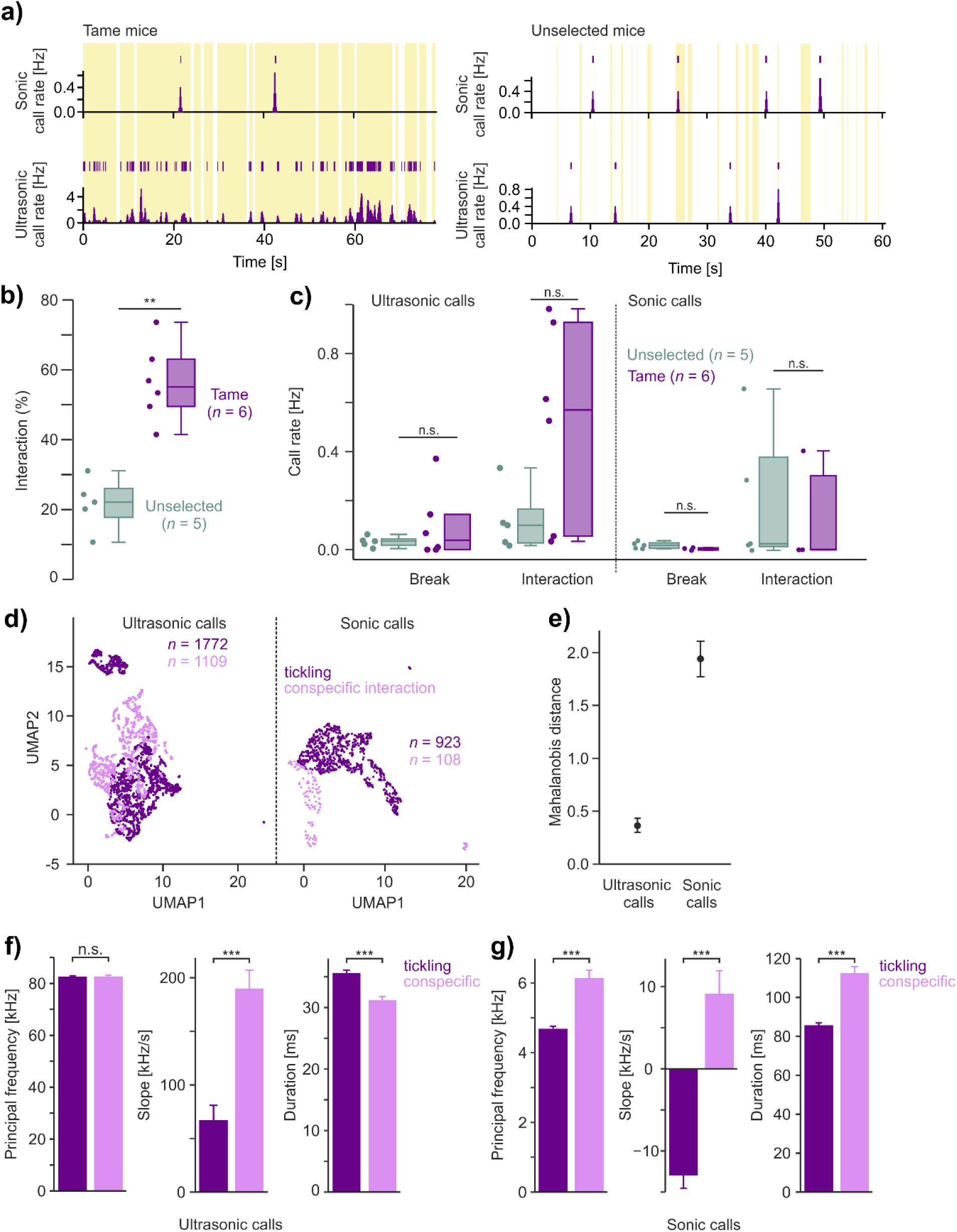
Behavioral characteristics during conspecific play in mice. **a)** Raster plot and histogram of sonic and ultrasonic call rate in a representative conspecific interaction session of a tame mouse pair (left) and an unselected mouse pair (right). Yellow boxes: interactions. **b)** Interaction percentage during conspecific interaction sessions. *n*: number of sessions; rank-sum test. **c)** Ultrasonic and sonic vocalization rate during different conspecific session events. *n*: number of sessions; rank-sum test. **d)** Projection of UMAP1 and 2 of calls features during tickling (purple) and conspecific interaction (lilac) sessions. *n*: number of calls. **e)** Mahalanobis distance (mean & 95% CIs) between tickling and conspecific interaction calls. **f)** Comparison of ultrasonic call features during tickling and conspecific interactions; rank-sum tes8t. **g)** Same as **f)** but for sonic calls. All *p*-values are detailed in Supplementary Table 2.

There was no significant difference in either ultrasonic or sonic call rates between the two groups during both interaction and break periods (Figure 2c). However, it is worth noting that the ultrasonic call rate for tame mice during the interaction period appeared higher, though not statistically significant, possibly due to little data and individual variations among tame mice (0.57 [0.055, 0.927] vs. 0.097 [0.025, 0.160]; median & IQR; *p*-value = 0.1).

To explore whether heterospecific play with humans and conspecific play share similar characteristics in tame mice, call features were projected in UMAP space for both ultrasonic and sonic calls recorded during both types of interactions (Figure 2d; Supplementary Figure 1). The results revealed a distinct separation of ensembles in both conditions for each call type, indicating different features of calls during human and conspecific interactions. This distinction was further quantified by calculating the Mahalanobis distance between tickling-induced and conspecific interaction-induced calls for each call type (Figure 2e). The positive confidence intervals for both ultrasonic and sonic calls signified substantial differences during each context. Differences in ultrasonic call characteristics, particularly slope and duration, were evident when comparing tickling to conspecific interactions, whereas principal frequency remained consistent (Figure 2f). Conversely, significant differences were observed across all these features for sonic calls (Figure 2g). This suggests that tame mice demonstrate distinct vocalization patterns during conspecific interactions, emphasizing context-dependent nature of play behavior with conspecifics and its differentiation from human interaction.

### Mice show no change in general anxiety level across days and between conditions

General anxiety in mice was assessed over several days, examining variables such as time spent in the center and total distance travelled. However, no significant changes were observed across days within each breeding group, in addition to no differences between the groups (Supplementary Figure 2).

### Hamsters minimally play with humans while engaging in conspecific play with context-dependent vocalizations

We observed hamsters vocalizing throughout tickling sessions (Figure 3a & Supplementary Video 6). However, call emissions were not clearly modulated by tactile stimuli (Figure 3b). During the introduction period, few vocalizations were observed (0.02 [0.00, 0.03] Hz; median & IQR). Call rates were higher during dorsal tickling (0.05 [0.02, 0.10] Hz; median & IQR) than during introduction (*p*-value = 0.002), but remained stable during breaks (0.04 [0.01, 0.10] Hz; median & IQR; *p*-value = 0.90). Call rates during dorsal gentle touch (0.02 [0.00, 0.06] Hz; median & IQR), on the other hand, were lower than breaks (*p*-value = 0.014). Hamsters moderately chased experimenters’ hand during no-contact play phases (Figure 3c). On the contrary, when paired, hamsters exhibited robust interactions with notable call emissions (Figure 3d & Supplementary Video 7), occurring more frequently during interaction periods compared to breaks (Figure 3e). Additionally, in some conspecific sessions, we observed audible squeaks at a very low rate at around 0.01 Hz in 5 out of 11 sessions, with one session having higher squeak rate at 0.11 Hz. Overall, hamsters engaged extensively in conspecific interactions, with interaction rates averaging approximately 50% of the session time (Figure 3f). Comparing call emissions between tickling and conspecific contexts revealed significantly higher vocalizations during conspecific interactions (Figure 3g).

**Figure 3:**
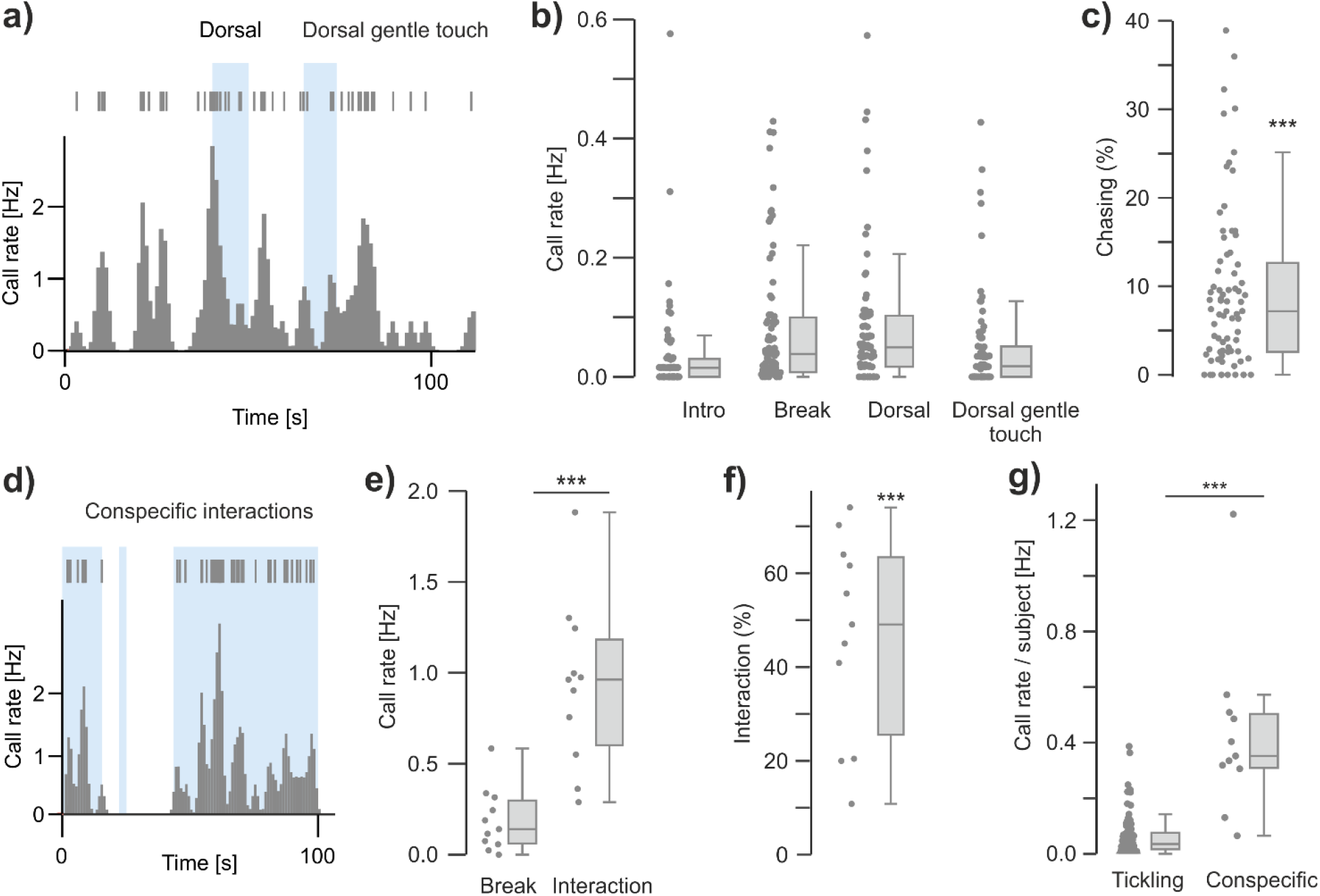
Behavioral characteristics during human-hamster and hamster-hamster play interactions. **a)** Raster plot and histogram of call rate in a representative tickling session. Blue boxes: interaction events. **b)** Call rate during different heterospecific interaction events. **c)** Chasing time percentage during no-contact play phases; one-sample t-test. **d)** Raster plot and histogram of call rate in a representative conspecific interaction session. Blue boxes: interaction events. **e)** Call rate during different heterospecific interaction and breaks; Wilcoxon signed-rank test. **f)** Interaction time percentage during conspecific interaction sessions; one-sample t-test. **g)** Call rate per subject during heterospecific and conspecific interaction events; rank-sum test. All *p*-values are detailed in Supplementary Table 2.

To assess whether the contextual differences in tickling and conspecific interactions manifest in call features, and given the unclear distinction between ultrasonic and sonic calls in hamsters compared to mice, we employed spectrogram image embedding into UMAP space [37] (Supplementary Figure 1). This analysis delineated clear separation between tickling and conspecific clusters (Figure 4a), with conspecific interaction calls forming a single cluster compared to the five clusters observed during tickling including an outlier cluster i.e. Cluster 0 (Figure 4b).

**Figure 4:**
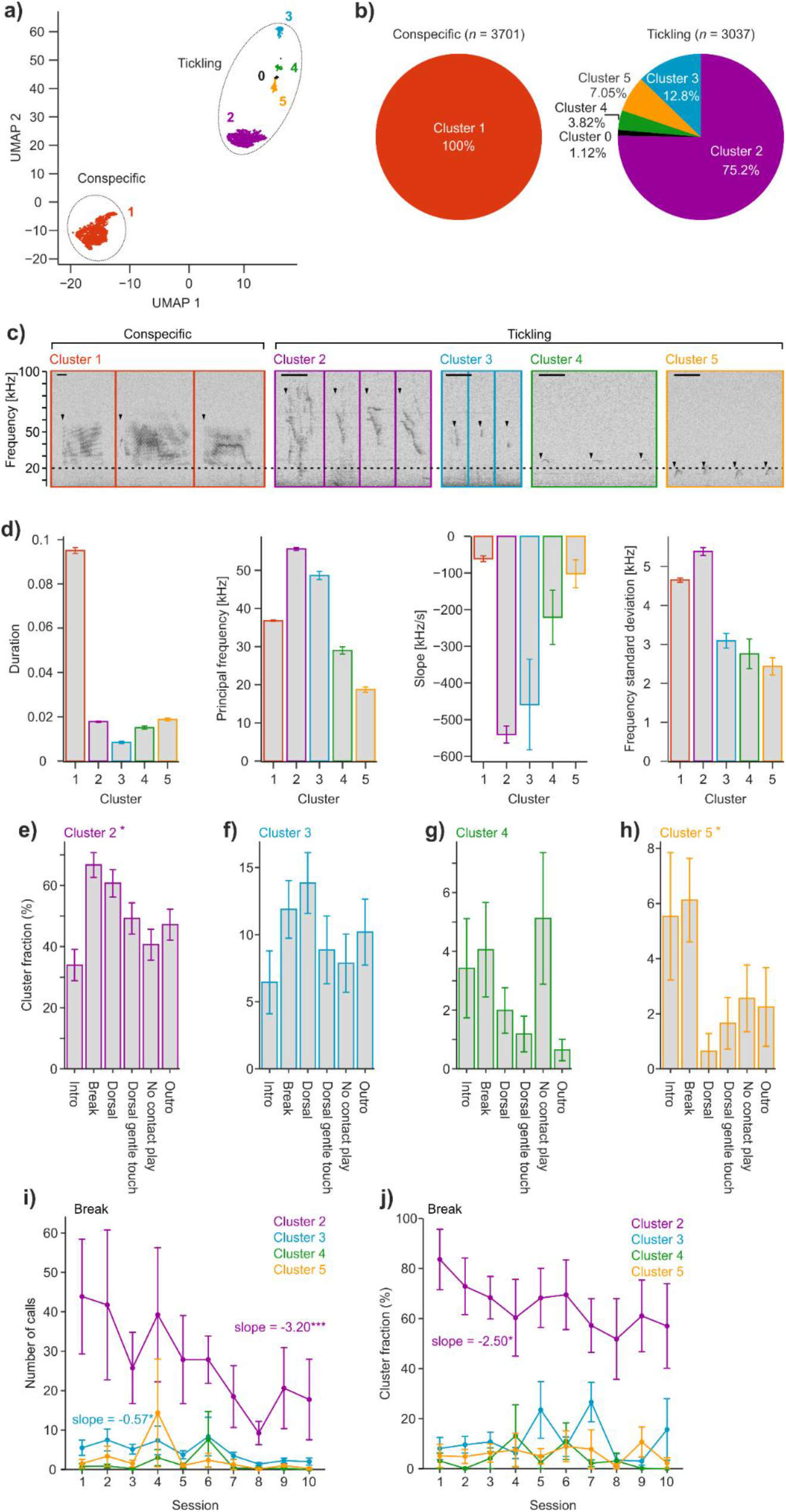
Context-dependent vocal patterns in hamsters. **a)** Projection of UMAP1 and 2 of spectrogram images during tickling and conspecific interaction sessions. Clustering was performed using HDBSCAN. **b)** Call cluster distributions for tickling and conspecific sessions. *n*: number of calls. **c)** Representative spectrograms for each call cluster during conspecific and tickling sessions. Arrow heads: call start time; scale bar: 50 ms. Dashed line indicates 20 kHz human hearing limit. **d)** Comparison of call features between clusters; rank-sum test. **e)** Fraction of cluster 2 calls during different events. Mean ± SEM; Kruskal-Wallis test; n = 78 sessions. **f-h)** Sam1e1as **e)** but for cluster 3-5 calls. **i)** Number of calls for each call clusters over tickling sessions. Mean ± SEM; n = 8 subjects; slope: linear mixed-effect model. **j)** Fraction of clusters over tickling sessions. Mean ± SEM; n = 8 subjects; slope: linear mixed-effect model. All *p*-values are detailed in Supplementary Table 2.

Calls emitted during conspecific interactions exhibited uniform ultrasonic characteristics, featuring long duration, flat slope, and complex structures, comprising parallel evenly-spaced lines representing multiples of the fundamental frequency i.e. harmonics, overlaid by dynamic, stroke-like trajectories, i.e. formants (Figure 4c, d & Supplementary Video 7). These calls resemble vocalizations previously described in the sexual context in hamsters [39]. Conversely, tickling-induced calls displayed greater variability, with distinct ultrasonic characteristics such as high principal frequency and sharp negative slope (Cluster 2), short ultrasonic calls (Cluster 3), and inverted-U shapes, which were ultrasonic in Cluster 4 but sonic in Cluster 5 (Figure 4c, d & Supplementary Video 6). Further analysis revealed that the clusters obtained varied in proportions across events, emphasizing the context dependency of call emission in Cluster 2 and 5 (Figure 4e-h). For instance, Cluster 2, which comprised approximately 30% during intro, doubled to around 70% during breaks (Figure 4e). Mixed effects model analysis revealed a significant negative trend in the Cluster 2 fraction over recording sessions (slope = -2.50, p = 0.045), indicating a decrease of Cluster 2 calls by 2.50% per session, after controlling for random effects due to individual differences (Figure 4i, j).

### Hamsters spend more time in the center of the anxiety assessment arena across days

General anxiety in hamsters was assessed over several days, focusing on the time spent in the center and total distance travelled. Across experiment days, there was an increase in the time that hamsters spent in the center of the arena but not in the total distance travelled (Supplementary Figure 3b, e).

## Discussion

In this study, we analysed vocal and behavioral response to tickling and conspecific interactions in tame mice and hamsters. So far, only rats and some non-human primate species were reported to respond playfully to tickling by humans with appetitive vocalizations [22, 23] despite abundant anecdotal observations of human-pet playful interactions. Notably, it has been acknowledged in rodent vocalization community that, unlike rats, standard laboratory mice do not respond to tickling, despite the lack of publication endorsing this fact. We addressed this gap in literature driven by the curiosity of what makes rats but not mice play with humans. Thus, we tickled tame mice, a selectively bred line chosen for increased human interaction, and unselected control mice [30, 40], as well as hamsters, a solitary but popular species kept as pets. Furthermore, we compared vocal response during conspecific interactions in these species.

Tame mice showed increased playfulness with humans assessed by increased ultrasonic vocalizations and pronounced chasing behavior compared to unselected control mice (Figure 1f, i), while sonic calls during tickling interactions were not affected (Figure 1e). These results suggest that ultrasonic but not sonic vocalizations may represent playfulness towards humans, implying distinct functional roles within the vocal repertoire. Interestingly, tame mice showed an increase also in conspecific interactions (Figure 2b) without any alterations in both ultrasonic and sonic call rates (Figure 2c). It is, however, possible that further data acquisition might influence the observed results. We previously reported similar ultrasonic call rates between tame and unselected mice when exposed to an anesthetized female [32]. The profound impact of selective breeding on both human-mouse and conspecific interactions among tame mice pairs suggests the existence of a common underlying genetic and neurobiological mechanism governing sociability. On the other hand, features of ultrasonic and sonic vocalizations emitted during tickling and conspecific interactions led to a clear separation of calls within each context in tame mice (Figure 2d-g). This suggests that tame mice emit distinct vocalizations depending on the specific interaction context. The significance of this differentiation lies in the understanding that tame mice, despite exhibiting playfulness and social approach in both scenarios, modulate their vocal communication in a context-dependent manner. Consistent with previous results [32], general anxiety levels in the open field test did not differ between tame and unselected mice (Supplementary Figure 2), indicating that tameness and anxiety may have distinct molecular and genetic underpinnings.

Hamsters, on the other hand, showed little if not no overall engagement in playful interactions with humans, reflected in minimal chasing behavior of less than 10% on average (Figure 3c). In contrast, tame mice exhibited significant chasing behavior, accounting for approximately 50% of the event time (Figure 1i), while unselected control mice showed around 30% chasing time. It is important to note that longer-term observation might have revealed shifts in behavior or the emergence of additional traits. Rats, on the other hand, as observed in our unpublished data, engage in chasing behavior in similar contexts at a rate of around 75%. Earlier advances on hamster vocalization focused on isolation calls in pups [41], as well as sexual and aggressive calls in adults [39, 42]. While behavioral repertoire of social play has been extensively studied [33, 43, 44], little is known about vocalizations during play behavior in hamsters. In our study, hamsters vocalized throughout tickling sessions, but the vocal emissions were not clearly modulated by tickling stimuli (Figure 3b). It could be speculated that these calls signify aversiveness, although similar types of calls emitted during human interaction, to our knowledge, have been reported in pup hamsters [45] but not adults. In particular, the reduction of Cluster 2 calls, which occupied 75% of all calls during tickling sessions, across days (Figure 4 i-j) and the increase in time in center in the open field test (Supplementary Figure 3) suggest a potential association of this specific call type and general anxiety. Other call types might serve another social function, or perhaps they stem from investigation or curiosity. It would be of interest the explore the potential communicatory roles of these vocalizations through playback studies. On the other hand, hamsters highly engaged in complex conspecific interactions and rough and tumble play as previously shown [33]. These interactions were associated with an increase in call rate (Figure 3d-f) in line with earlier reports showing the emission of calls in different social contexts in hamsters [39, 46]. The rare occurrence of squeaks observed in both tickling and conspecific sessions are likely linked to aggression, as previously indicated [46]. Moreover, the variability in squeak occurrence across conspecific sessions may be attributable to factors such as hierarchy or compatibility among playmates. Spectrogram analysis revealed homogeneous calls emitted during conspecific interactions compared to highly heterogeneous calls emitted during tickling (Figure 4a-d), highlighting the context-specific vocal responses similar to tame mice.

Human-animal interactions in particular play behavior have been well documented typically in domesticated species such as dogs and cats [1, 19-21, 47, 48] as well as rats [49]. However, the biological basis of human-directed social engagement remains elusive. Selective breeding for tameness, i.e. affinity towards humans, in non-playful wild mice stock led to a high level of social human engagement and receptiveness after only 36 generations. In addition to the upregulated oxytocin receptor gene in tame mice [31], the possible associated genes within the genomic regions active tameness region 1 (ATR1) and active tameness region 2 (ATR2) on chromosome 11, which are shared with domesticated dogs [31], may potentially indicate a genetic foundation for the observed playfulness. Given that selective breeding for tameness not only increased playfulness towards humans, but also within conspecifics, it is plausible to propose that these genes may play a role in fostering social play across various contexts, irrespective of the playmate involved. Notably, several of these genes, including *ATP5G1* and *Sez6*, have established associations with neuropsychiatric disorders such as schizophrenia, bipolar disorder, and depression, all of which are characterized by social deficits [50-52]. Furthermore, the *Srr* gene has been implicated in causing a reduction in dopamine release in the ventral tegmental area in *Srr*-knockout mice following METH administration [53]. While these specific genomic areas have not been directly associated with social behavior or playfulness, the correlation supports the hypothesis that certain genetic factors may influence the behavioral tendencies in tame mice. Could the shared genetic elements between tame mice and domesticated dogs be indicative of a broader phenomenon, one that extends to various rodent species and aligns with the concept of self-domestication? This raises intriguing questions about the interplay of genetics, behavior, and self-domestication and invites a comparative analysis with other species.

The domestication of animals, for instance dogs and cats, has historically been influenced by considerations of utility, companionship, and desirable behavioral traits. In contrast to dogs, selected for roles like hunting and guarding, and cats, valued for pest control, mice lacked attributes making them initially appealing for domestication. Their small size and prey status may have limited their immediate utility to early human societies. Furthermore, inherent traits, such as an aversion to human touch and specific environmental adaptations, could have hindered their domestication compared to more adaptable species. The nuanced dynamics of mutual benefit, often integral to domestication processes, may not have aligned with human needs or preferences in the historical context of mice. Golden hamsters, on the other hand, did not show an inclination towards forming playful bonds with humans (Figure 3). Social behaviors and play interactions within their species has been extensively reported [43, 54] and may serve purposes such as communication, hierarchy establishment, or mate selection in addition to coping with stress [8, 9, 43]. This stands in contrast to the lack of similar behavior towards humans, indicating a distinction in their social cognition. This might be influenced by their evolutionary history as nocturnal rodents with a solitary lifestyle, potential genetic predispositions, and the initial domestication for research purposes rather than direct human companionship [55].

Tame mice and rats, which are more social and communal rodents, on the other hand, show a remarkable inclination toward human interaction and playfulness. Indeed, rats are known for their adaptability and social behaviors including response to tickling and human approach [23, 24]. While it can be tempting to suggest that rats are associated with disease because of the bubonic plague, human disgust with rats is neither ahistorical nor innate. In Europe, rats were not even suspected vectors of the plague until hundreds of years after its devastation [56]. The ways in which humans view animals in general draws from cultural connotations [57]. Rats have flourished with human health population events like the rise of cities. With the rise of cities, large populations of people needed to manage waste removal at a large scale that could not be easily integrated back into the environment. This was also the point in time when humans began to differentiate rats from mice, a distinction which did not exist before the 17th century [58]. Norway rats were able to displace ship rats, which look a bit more like mice. Humans saw Norway rats flourish in sewers and the trash, leading to a new name for them, sewer rats. Sewer rats who learn how to navigate human infrastructures and human schedules survived the best, perhaps leading to their tendency toward self-domestication, and, as a generalist species like humans, they adapted to nearly all places humans inhabit.

Vocalizations have been extensively studied across various rodent species such as mice, hamsters, and rats [24, 39, 59-62], focusing on their categorization with a large degree of consensus among scientists regarding concepts like appetitive versus aversive calls. However, much remains unknown, particularly regarding the context-specific nature of vocal patterns or syllables. This idea finds parallels in studies of birds, where specific sequences have been identified for distinct communicative purposes [63, 64] as well as other describing differences in calls of rats between tickling and conspecific play [65]. Our results also reflect this concept, as the vocalizations of mice and hamsters displayed distinct patterns determined by both the context of interaction and the species of their playmates (Figure 2&4). The implications of such context-specific responses highlight the remarkable cognitive abilities of both mice and hamsters, namely their social intelligence that surpasses current understandings of rodent vocalizations and social perception.

In conclusion, this study unravelled the interplay of selective breeding and context-dependent vocalizations in rodents. Tame mice, specifically bred for human interaction, exhibited playful behavior towards both humans and conspecifics, solidifying their potential as models for understanding human-animal bonds. While hamsters showed minimal playful interaction with humans, both tame mice and hamsters displayed unique vocal repertoires depending on the social context, suggesting species-specific social preferences and cognitive abilities to adapt their response to different social scenarios. Further exploration of the genetic basis underlying playfulness in tame mice could yield valuable insights into human-animal bonds and potentially inform our understanding on neuropsychiatric disorders. Overall, this research opens exciting avenues to delve deeper into the intricate relationship between genetics, behavior, and self-domestication in rodents, ultimately illuminating human-animal interactions and the evolution of social behavior across diverse species.

## Supplementary Materials

Supplementary Videos 1-7

Supplementary Figures 1-3

Supplementary Tables 1-2

## Acknowledgments and Funding

Authors thank Jannik Wagner for assisting software development, Nicholas Jourjine for comments on hamster vocalizations, Juan Ignacio Sanguinetti-Scheck for discussion, and Eduard Maier for comments on manuscript. This study was supported by ‘Freigeist’ Fellowship (VolkswagenStiftung) to S.I., JST PRESTO (JPMJPR2048) to H.N., and the Grant-in-Aid for Scientific Research (JSPS KAKENHI Grant Number 19KK0177) to T.K.

## Author contributions

Conceptualization, S.D., H.N., T.K. and S.I.; Experiments, S.D. and S.I., Formal analysis, S.D. and S.I.; Visualization, S.D.; Funding acquisition, H.N., T.K., and S.I.; Supervision, H.N., T.K., and S.I.; Writing-original draft, S.D., D.D. and S.I.; Technical supervision on hamster handling: R.Y.S. Comments and draft editing: S.D., H.N., T.K., and S.I.

