## Supplementary file for "Comparative Analysis of Tickling and Conspecific Play in Tame Mice and Golden Hamsters"

**Supplementary Table 1: Behavioral ethograms in video analysis labelling**

| **Experiment** | **Ethogram** | **Description** |
|---|---|---|
| Tickling | Dorsal | Tickling on the dorsal trunk |
| | Dorsal gentle touch | Gentle stroking on the dorsal trunk |
| | No-contact play | Experimenter’s hand in the arena without physical contact with the animal |
| | Chasing | Animal chasing the experimenter’s hand |
| | Vertical jump (mice) | Mouse vertical jumping behavior |
| | Intro | First 2 min without interaction |
| | Break | Periods between interactions |
| | Outro | Last 2 min without interaction |
| Conspecific interaction | Interaction | Physical contact of the two animals |
| | Break | Periods without physical contact of the two animals |

**Supplementary Table 2: Statistical *p*-values**

| Figure | Panel | Variable | test | *p*-value |
|---|---|---|---|---|
| 1 | f | Dorsal USVs | rank-sum | < 0.001 |
| 1 | f | Dorsal gentle touch USVs | rank-sum | 0.070 |
| 1 | f | No contact play USVs | rank-sum | 0.630 |
| 1 | f | Dorsal sonic | rank-sum | 0.070 |
| 1 | f | Dorsal gentle touch sonic | rank-sum | 0.160 |
| 1 | f | No contact play sonic | rank-sum | 0.850 |
| 1 | g | Jump rate | rank-sum | 0.020 |
| 1 | h | Probability | rank-sum | < 0.001 |
| 1 | i | Chasing (%) | rank-sum | 0.005 |
| 2 | b | Interaction (%) | rank-sum | 0.006 |
| 2 | c | Break USVs | rank-sum | 0.860 |
| 2 | c | Interaction USVs | rank-sum | 0.100 |
| 2 | c | Break sonic | rank-sum | 0.180 |
| 2 | c | Interaction sonic | rank-sum | 0.650 |
| 2 | f | Principal frequency USVs | rank-sum | 0.160 |
| 2 | f | Slope USVs | rank-sum | < 0.001 |
| 2 | f | Duration USVs | rank-sum | < 0.001 |
| 2 | f | Principal frequency sonic | rank-sum | < 0.001 |
| 2 | f | Slope sonic | rank-sum | < 0.001 |
| 2 | f | Duration sonic | rank-sum | < 0.001 |
| 3 | b | Intro vs Break | Linear mixed model | < 0.001 |
| 3 | b | Intro vs Dorsal | Linear mixed model | 0.002 |
| 3 | b | Intro vs Dorsal gentle touch | Linear mixed model | 0.170 |
| 3 | b | Break vs Dorsal | Linear mixed model | 0.900 |
| 3 | b | Break vs Dorsal gentle touch | Linear mixed model | 0.014 |
| 3 | b | Dorsal vs Dorsal gentle touch | Linear mixed model | 0.047 |
| 3 | c | Chasing (%) | One sample t-test | < 0.001 |
| 3 | e | Call rate | Wilcoxon signed-rank | < 0.001 |
| 3 | f | Interaction (%) | One sample t-test | < 0.001 |
| 3 | g | Call rate/subject | rank-sum | < 0.001 |
| 4 | d | Duration 1 vs 2 | rank-sum | < 0.001 |
| 4 | d | Duration 1 vs 3 | rank-sum | < 0.001 |
| 4 | d | Duration 1 vs 4 | rank-sum | < 0.001 |
| 4 | d | Duration 1 vs 5 | rank-sum | < 0.001 |
| 4 | d | Duration 2 vs 3 | rank-sum | < 0.001 |
| 4 | d | Duration 2 vs 4 | rank-sum | 0.100 |
| 4 | d | Duration 2 vs 5 | rank-sum | < 0.001 |
| 4 | d | Duration 3 vs 4 | rank-sum | < 0.001 |
| 4 | d | Duration 3 vs 5 | rank-sum | < 0.001 |
| 4 | d | Duration 4 vs 5 | rank-sum | < 0.001 |
| 4 | d | Principal frequency 1 vs 2 | rank-sum | < 0.001 |
| 4 | d | Principal frequency 1 vs 3 | rank-sum | < 0.001 |
| 4 | d | Principal frequency 1 vs 4 | rank-sum | < 0.001 |
| 4 | d | Principal frequency 1 vs 5 | rank-sum | < 0.001 |

| | | | | |
|---|---|---|---|---|
| 4 | d | Principal frequency 2 vs 3 | rank-sum | < 0.001 |
| 4 | d | Principal frequency 2 vs 4 | rank-sum | < 0.001 |
| 4 | d | Principal frequency 2 vs 5 | rank-sum | < 0.001 |
| 4 | d | Principal frequency 3 vs 4 | rank-sum | < 0.001 |
| 4 | d | Principal frequency 3 vs 5 | rank-sum | < 0.001 |
| 4 | d | Principal frequency 4 vs 5 | rank-sum | < 0.001 |
| 4 | d | Slope 1 vs 2 | rank-sum | < 0.001 |
| 4 | d | Slope 1 vs 3 | rank-sum | < 0.001 |
| 4 | d | Slope 1 vs 4 | rank-sum | 0.004 |
| 4 | d | Slope 1 vs 5 | rank-sum | 0.036 |
| 4 | d | Slope 2 vs 3 | rank-sum | 0.020 |
| 4 | d | Slope 2 vs 4 | rank-sum | < 0.001 |
| 4 | d | Slope 2 vs 5 | rank-sum | < 0.001 |
| 4 | d | Slope 3 vs 4 | rank-sum | 0.060 |
| 4 | d | Slope 3 vs 5 | rank-sum | 0.001 |
| 4 | d | Slope 4 vs 5 | rank-sum | 0.390 |
| 4 | d | Frequency standard deviation 1 vs 2 | rank-sum | 0.100 |
| 4 | d | Frequency standard deviation 1 vs 3 | rank-sum | < 0.001 |
| 4 | d | Frequency standard deviation 1 vs 4 | rank-sum | < 0.001 |
| 4 | d | Frequency standard deviation 1 vs 5 | rank-sum | < 0.001 |
| 4 | d | Frequency standard deviation 2 vs 3 | rank-sum | < 0.001 |
| 4 | d | Frequency standard deviation 2 vs 4 | rank-sum | < 0.001 |
| 4 | d | Frequency standard deviation 2 vs 5 | rank-sum | < 0.001 |
| 4 | d | Frequency standard deviation 3 vs 4 | rank-sum | < 0.001 |
| 4 | d | Frequency standard deviation 3 vs 5 | rank-sum | < 0.001 |
| 4 | d | Frequency standard deviation 4 vs 5 | rank-sum | 0.840 |
| 4 | e | Cluster 2 | Kruskal-Wallis | 0.043 |
| 4 | f | Cluster 3 | Kruskal-Wallis | 0.493 |
| 4 | g | Cluster 4 | Kruskal-Wallis | 0.456 |
| 4 | h | Cluster 5 | Kruskal-Wallis | 0.014 |

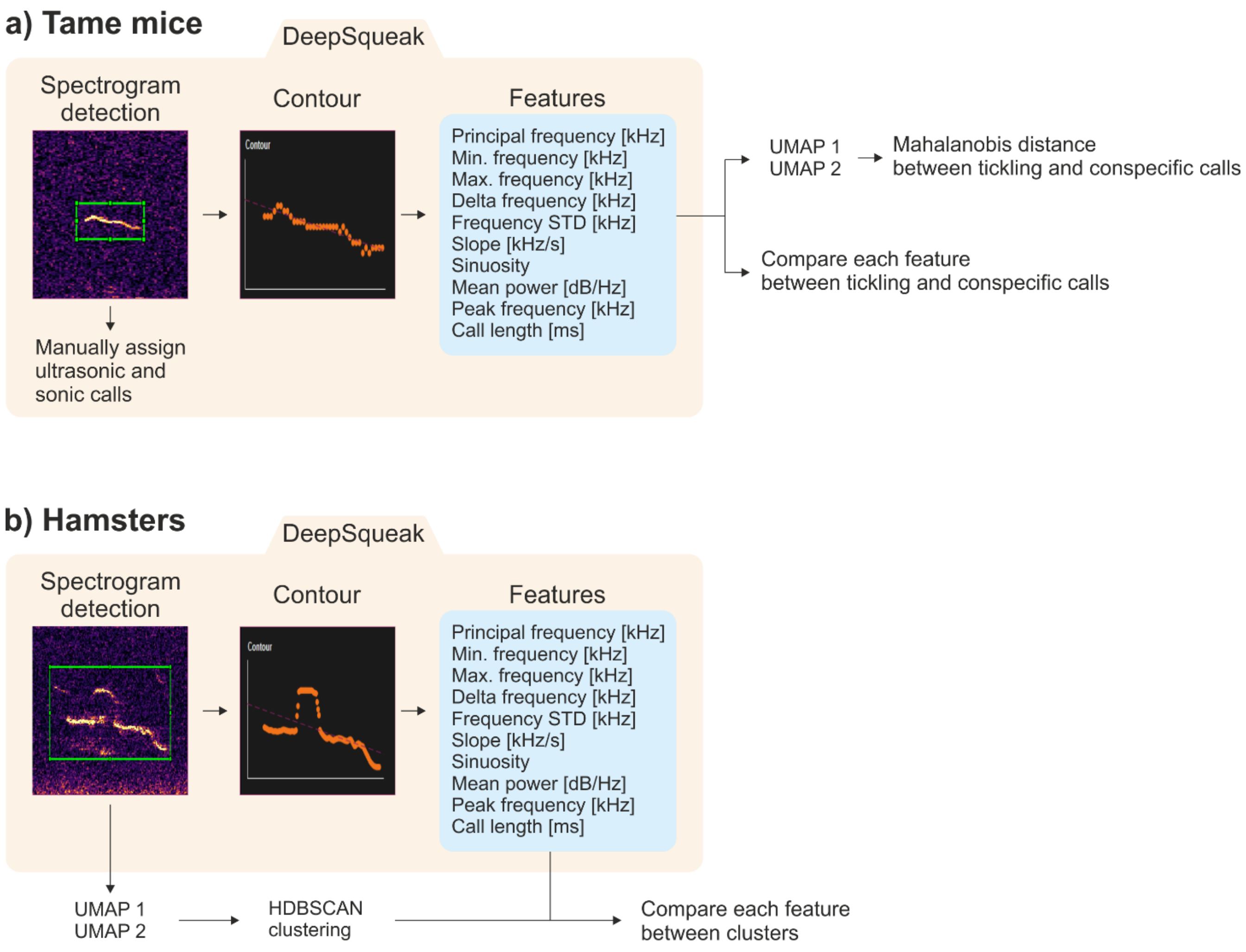


**Supplementary Figure 1: Call analysis workflow**
**a)** Call analysis for tame mice as detailed in *Materials and Methods*. **b)** Same as **a)** but for hamsters

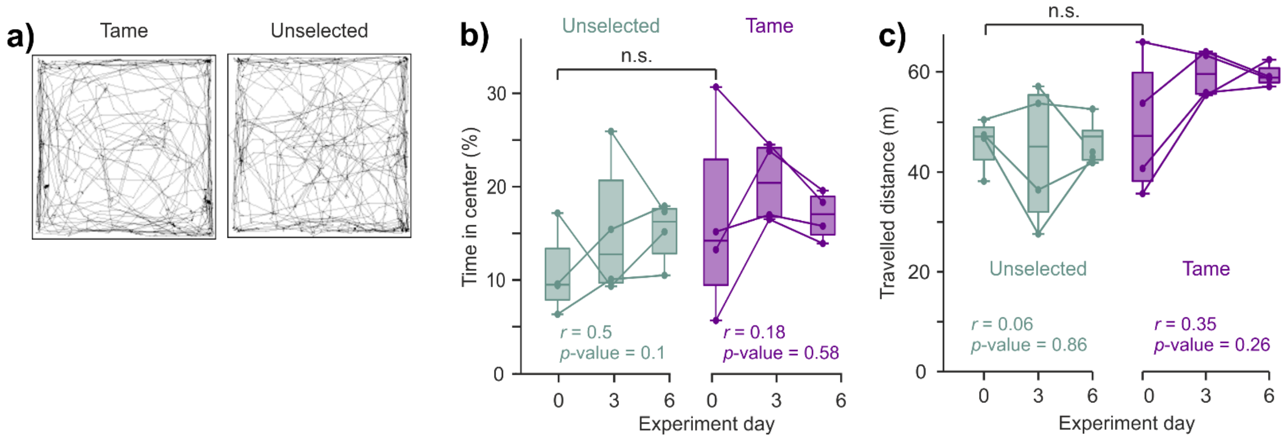


**Supplementary Figure 2: Anxiety assessment using open field test in mice**
**a)** Representative trajectories of tame and control unselected mice in open field chamber over 10 min. **b)** Percentage of the time spent in the center across test days for unselected controls mice (green) and tame mice (purple). Dots represent sessions; lines represent individual subjects; Baseline comparison on Day 0 between the groups was assessed using rank-sums test; *r* represents Spearman's rank correlation coefficient. **c)** Same as b) but for distance travelled.

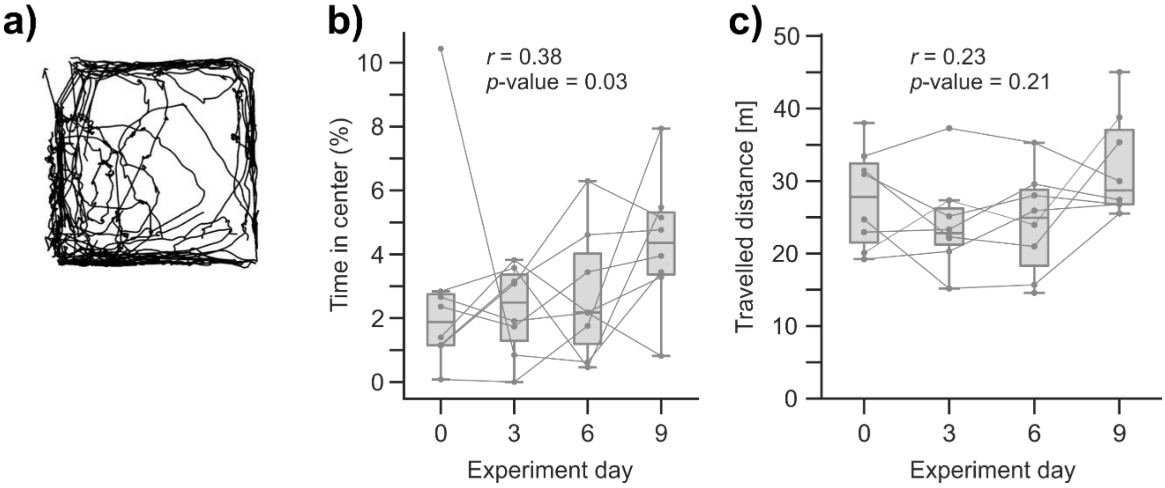


**Supplementary Figure 3: Anxiety assessment using open field test in hamsters**
**a)** Representative trajectories of hamsters in open field chamber over 10 min. **b)** Percentage of the time spent in the center across test days. Dots represent sessions; line represents individual subjects; *r* represents Spearman's rank correlation coefficient. **c)** Same as b) but for distance travelled.

**Supplementary Video 1:** Tame mouse tickling

**Supplementary Video 2:** Unselected mouse tickling

**Supplementary Video 3:** Unselected mouse jumps

**Supplementary Video 4:** Tame mouse chasing

**Supplementary Video 5:** Unselected mouse no-contact play

**Supplementary Video 6:** Hamster tickling

**Supplementary Video 7:** Hamsters conspecific play
